# Concreteness shapes semantic representations in bilingual brains

**DOI:** 10.64898/2025.12.23.696243

**Authors:** Mathis Lamarre, Xue L. Gong, Catherine Chen, Fatma Deniz

## Abstract

Behavioral studies show that humans process concrete words more similarly across languages than abstract words. This suggests that in bilingual brains, semantic representations may be more similar across languages for concrete concepts than for abstract concepts, but existing neuroimaging evidence is inconclusive. Here, we analyzed functional magnetic resonance imaging (fMRI) data from fluent Chinese-English bilinguals who read several hours of naturalistic narratives in both languages. We used encoding models to estimate voxelwise tuning towards concrete and abstract concepts in each language separately. We then quantified the similarity of cortical semantic representations across languages. First, we find that the cortical organization of concreteness tuning is consistent across languages. Second, semantic representations are similar across languages for both concrete-tuned voxels and abstract-tuned voxels. Third, we find that for abstract concepts, cross-language similarity of semantic representations may be driven by emotionality. Overall, these findings reveal how concreteness affects semantic representations in bilingual brains.

## 1 Introduction

Concreteness, the degree to which a concept refers to a perceptible entity, plays a key role in psycholinguistic models of bilingualism (Pavlenko 2009). In these models, semantic representations are more similar across languages for concrete words than for abstract words (de Groot 1992; Van Hell and De Groot 1998). For example, these models predict that the word “house” is represented more similarly across languages than the word “truth”. This is supported by behavioral evidence that translation times are faster and cross-language priming is stronger for concrete words than for abstract words (Basnight-Brown and Altarriba 2016; Kiran and Tuchtenhagen 2005; Tokowicz and Kroll 2007; Ferre et al. 2017; Schoonbaert et al. 2009). These behavioral studies suggest that, in the brains of bilinguals, semantic representations may be more similar across languages for concrete concepts than for abstract concepts. Alternatively, behavioral differences between concrete and abstract concepts could arise from some other cognitive process in the brain. Thus, it remains unclear how concreteness affects the cross-language similarity of semantic representations in bilingual brains.

A few neuroimaging studies have investigated the role of concreteness on the similarity of semantic representations across languages in bilingual brains (Li et al. 2021, 2023). They found greater cross-language similarity of semantic representations for abstract words in regions typically associated with language, and for concrete words in sensorimotor regions. However, these studies have two key limitations. First, they used controlled stimuli, presenting isolated nouns without any context. Other work has shown that cortical representations of concrete and abstract concepts are strongly context-dependent (Kewenig et al. 2023), and that naturalistic stimuli engage a much broader set of cortical regions than isolated words (Hamilton and Huth 2020; Deniz et al. 2023). Thus, experiments with controlled stimuli may not fully characterize cortical representations of concrete and abstract concepts. Second, they modelled concreteness as a binary category rather than a continuous dimension. This discrete categorization limits the granularity of the analysis and may hide patterns across levels of concreteness. Overall, existing neuroimaging studies of semantic representations in bilinguals are insufficient to determine the role of concreteness on cross-language similarity of semantic representations.

Monolingual neuroimaging studies have shown where in the brain concrete and abstract concepts are represented for individual languages. A large body of work showed that concrete concepts and abstract concepts are represented in distinct cortical regions (Bedny and Thompson-Schill 2006; Binder et al. 2009; Tang et al. 2021). Concrete concepts are represented close to their corresponding perceptual systems (Barsalou 2008; Martin 2016) while abstract concepts are represented mainly in regions typically associated with language (Sabsevitz et al. 2005; Noppeney et al. 2004) or emotion (Vigliocco et al. 2014). However, most of this research was carried in English. More than half of the studies in a large meta-analysis (Wang et al. 2010) were in English, and the remaining ones in Germanic or Romance languages, which are not typologically distant from English. We could only find one monolingual study in Chinese (Kansaku et al. 1998) but it was limited to areas in the occipital and temporal cortices. This raises the question of whether the cortical organization of concrete and abstract concepts generalizes to languages that are more typologically distant (Blasi et al. 2022).

Languages that are typologically distant from English may process concreteness differently. Prior studies have shown that reading concrete words is faster and more accurate than reading abstract words, an observation known as the concreteness effect (Jessen et al. 2000). The concreteness effect has been shown to be stronger in logographic languages (like Chinese) than in alphabetic languages (like English) (Shibahara et al. 2003). It was also shown that reading Chinese characters elicits brain response patterns more similar to picture perception than those observed during English word reading (Chee et al. 2000). Thus, comparing typologically distant languages such as English and Chinese provides a critical test for the comparison of the representations of concrete and abstract concepts, both in terms of cortical organization and cross-language similarity.

Here, we examine how concrete and abstract concepts are represented in the brains of Chinese-English bilinguals. We analyzed functional magnetic resonance imaging (fMRI) recordings collected while participants read narratives in both English and Chinese (Huth et al. 2016; De Heer et al. 2017; Deniz et al. 2019; Lamarre et al. 2022; Chen et al. 2024a; Negi et al. 2025). All participants were native Chinese speakers who are fluent in both languages. Voxelwise encoding models were used to map cortical semantic representations for each participant in each language separately (Wu et al. 2006; Naselaris et al. 2011; Nunez-Elizalde et al. 2019). The estimated semantic model weights were used to measure each voxel’s preference for concrete or abstract concepts, which we define as *concreteness tuning*. For each voxel, the estimated semantic model weights were used to measure the similarity of semantic representations across the two languages, which we refer to as *cross-language similarity*. These two metrics were used to investigate how concreteness affects the similarity of semantic representation across languages in bilingual brains.

## 2 Results

### 2.1 Overview

We compared cortical representations of concrete and abstract concepts across languages in six Chinese-English bilinguals. To do this, we first used voxelwise encoding models to estimate weights that reflect the semantic information represented in each voxel (Fig. 1-A). This was performed separately for each participant and in each language. We used a lexical word embedding space (fastText (Bojanowski et al. 2017; Joulin et al. 2018)) to operationalize the semantic information conveyed by each word, and we replicated our results with a contextual embedding space (multilingual BERT, or mBERT (Devlin et al. 2019); see results in Supplemental Information). Low-level sensory features were regressed out of brain responses prior to model estimation (see Methods for details). We then used the estimated semantic model weights in three ways. First, we identified the concreteness dimension. The concreteness dimension is a vector in the embedding space spanning from abstract to concrete concepts. We projected the estimated semantic model weights onto this concreteness dimension to determine the concreteness tuning of each voxel, which measures the preference for concrete or abstract concepts (Fig. 1-B). Second, we computed the Pearson correlation of estimated semantic model weights across English and Chinese to measure the cross-language similarity of semantic representations (Fig. 1-C). Together, concreteness tuning and cross-language similarity allowed us to examine how concreteness influences cortical organization of semantic representations within language as well as the similarity of semantic representations across languages. Third, we identified the emotionality dimension–a vector in the embedding space spanning from concepts with low to high emotional content. We projected the estimated semantic model weights onto this concreteness dimension to determine the emotionality tuning of each voxel.

**Fig. 1.**
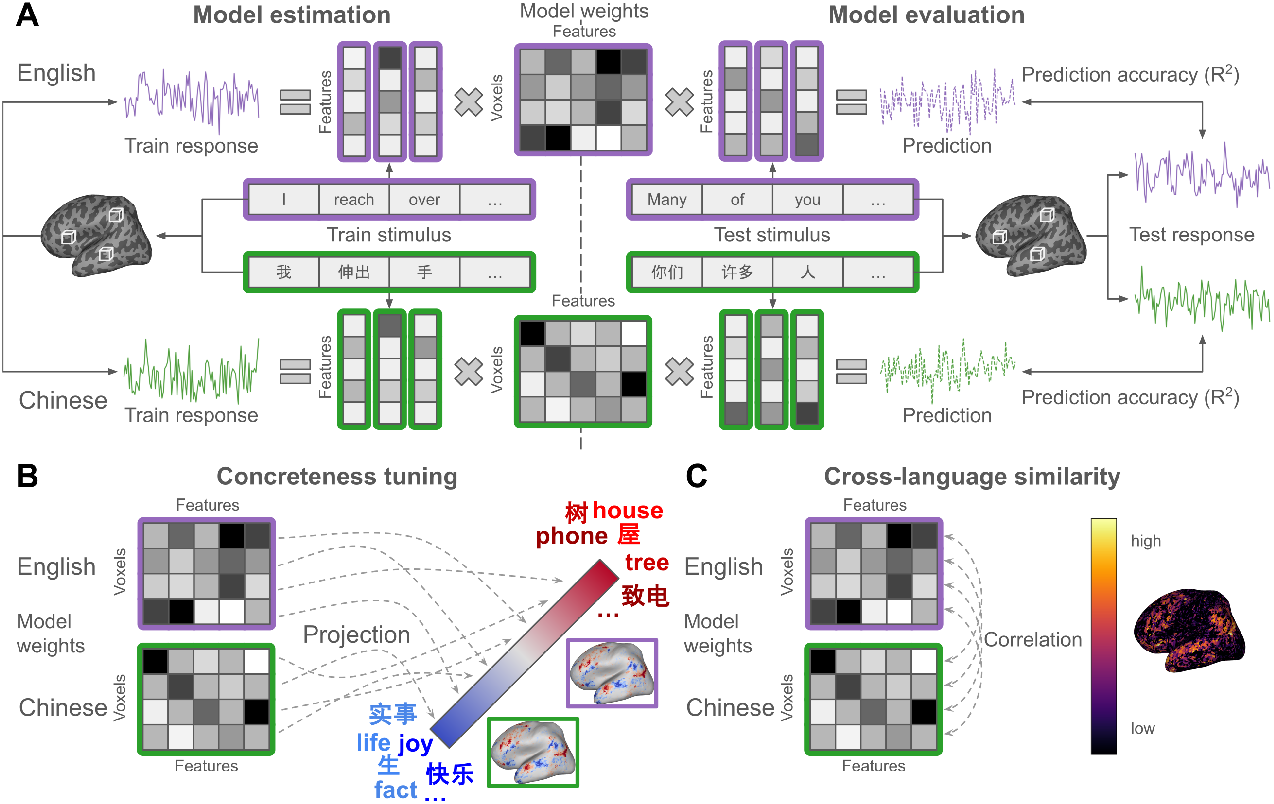
Voxelwise encoding model framework. **(A)** Experimental design and voxelwise encoding modeling. Brain responses were recorded using functional MRI while six Chinese-English bilinguals read the same narratives in both languages. Semantic features were constructed by extracting word embeddings for each word in the narratives. Model weights that describe the semantic representation of each voxel were estimated using regularized regression. Prediction accuracy was quantified as the coefficient of determination (*R*^2^) between predicted and recorded brain responses on a held-out narrative separately for each participant and language. **(B)** Measurement of concreteness tuning. The concreteness dimension is the vector in the word embedding space that spans from abstract to concrete concepts. The concreteness dimension was determined by taking the difference between the mean embedding of words rated as highly concrete and the mean embedding of words rated as highly abstract. Concreteness tuning was measured separately for each voxel and language as the projection of estimated semantic model weights onto the concreteness dimension. **(C)** Measurement of cross-language similarity. Cross-language similarity was measured for each voxel as the Pearson correlation between the estimated semantic model weights in English and Chinese.

With this metric, we evaluated the role of emotional content in the representations of concrete and abstract concepts and how it relates to cross-language similarity.

### 2.2 The cortical organization of concreteness tuning is consistent across languages

To measure the preference of each voxel for concrete or abstract concepts, we computed a concreteness tuning metric. To compute this metric, we first identified the concreteness dimension as a vector in the word embedding space that spans from abstract to concrete concepts. This is a necessary step because dimensions in the word embedding space do not have explicit meaning. To define the concreteness dimension, we identified words that humans rated as highly concrete and highly abstract in each language (Brysbaert et al. 2014; Xu and Li 2020). Next, for each language separately, we computed the concreteness dimension as the difference between the mean embedding of the concrete words and the mean embedding of the abstract words. To avoid bias towards the embedding space of either language, we then computed the average of the English and Chinese concreteness dimensions as the final concreteness dimension. We validated the concreteness dimension by comparing the human concreteness rating of each stimulus word with the projection of the word’s embedding onto the concreteness dimension. We found that the ratings and projections are strongly positively correlated (*r* = .77, *p <* .001 in English, *r* = .70, *p <* .001 in Chinese, by a two-sided t-test, Fig. S1-C). The correlation is also high when mBERT is used as the word embedding space (*r* = .71, *p <* .001 in English, *r* = .76, *p <* .001 in Chinese, by a two-sided t-test, Fig. S9-C). These results show that the concreteness dimension captures human judgements of concreteness.

Next, to determine whether the cortical organization of concreteness tuning differs between languages, we examined concreteness tuning in each language separately. Concreteness tuning is the projection of a voxel’s estimated semantic model weights onto the concreteness dimension. Fig. 2-A shows concreteness tuning across the cortical surface of one representative participant. Results are shown in English and in Chinese separately. The color of the voxels reflects their concreteness tuning ranging from abstract (blue) to concrete (red). Voxels that are poorly predicted by that language’s semantic model are shown in gray (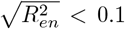 or 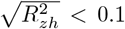). Our results show that, in both languages and bilaterally, posterior areas in the lateral and medial parietal cortices are tuned towards concrete concepts (voxels shown in red). In both languages and bilaterally, regions typically associated with language such as Broca’s area, the high-level auditory cortex, and the superior temporal sulcus are tuned towards abstract concepts (voxels shown in blue). The dorsolateral prefrontal cortex is tuned towards both concrete and abstract concepts. Fig. 2-B directly compares concreteness tuning between English (x-axis) and Chinese (y-axis) for the same representative participant. Each point reflects one voxel that is well predicted in at least one language. Concreteness tuning is strongly correlated between English and Chinese (*r* = .83, *p <* .001 by a two-sided t-test). The cortical organization of concreteness tuning is also consistent across participants (Fig. 2-C). In addition, the correlation between concreteness tuning in English and in Chinese is high for all participants (*r* = .82, .82, .85, .83, .89, *p <* .001 by a two-sided t-test for P2-6, Fig. S4).

**Fig. 2.**
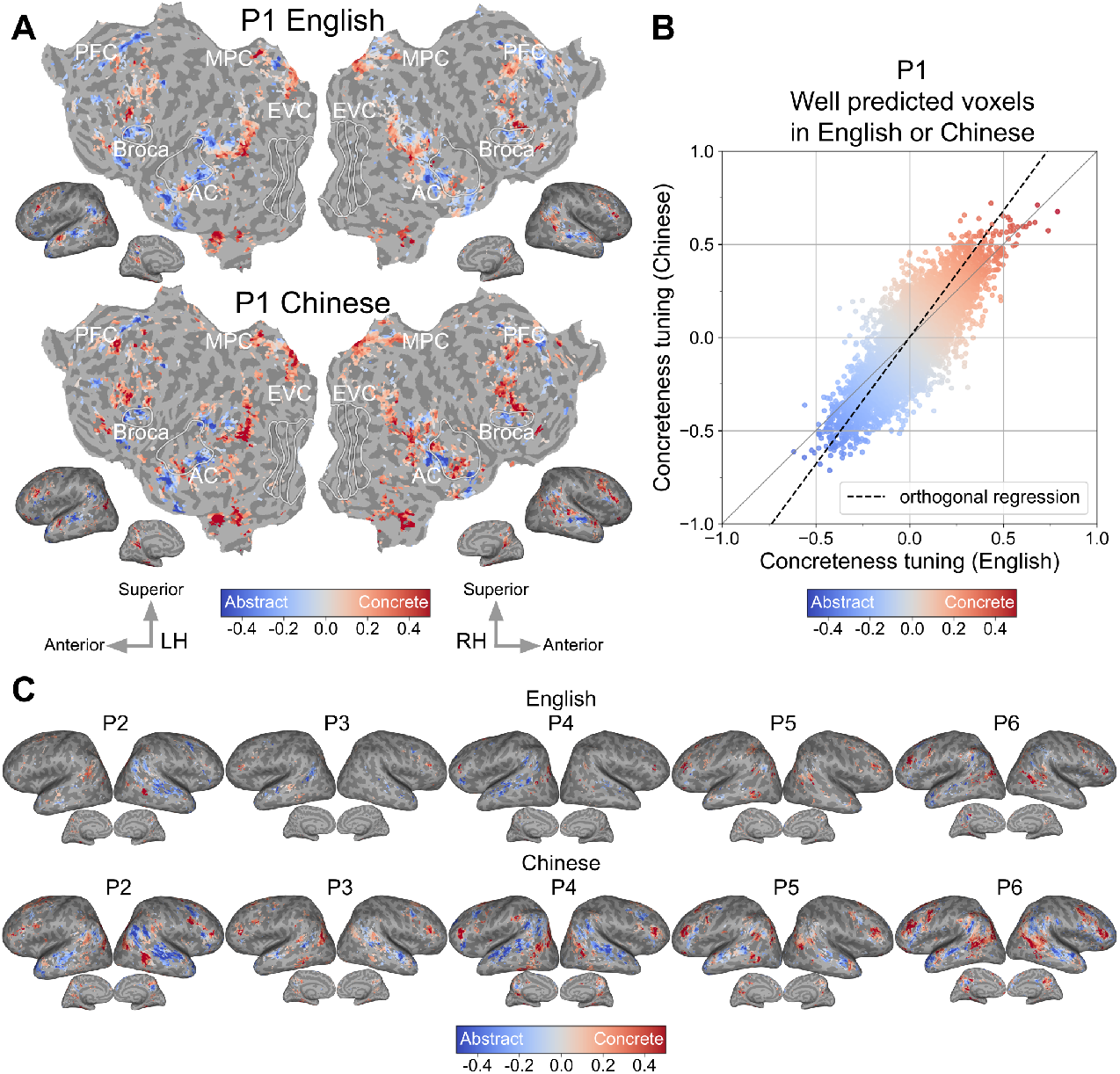
Concreteness tuning in English and Chinese. Concreteness tuning was measured in each language as the projection of each voxel’s estimated semantic model weights onto the concreteness dimension. The concreteness dimension is a vector in the word embedding space that spans from abstract to concrete concepts. **(A)** Concreteness tuning for English (top) and Chinese (bottom) is shown for one representative participant (P1) on their cortical surface. Voxel color indicates the value of concreteness tuning ranging from abstract (blue) to concrete (red). Voxels that were poorly predicted by the semantic model are shown in gray 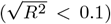. In both languages and bilaterally, posterior areas in the lateral and medial parietal cortices are tuned towards concrete concepts. Regions typically associated with language such as Broca’s area, the high-level auditory cortex and the superior temporal sulcus are tuned towards abstract concepts. The dorsolateral prefrontal cortex is tuned towards both concrete and abstract concepts. This suggests that the cortical organization of concreteness tuning is consistent across English and Chinese. **(B)** Comparison of concreteness tuning in English (x-axis) and Chinese (y-axis) in a scatterplot for the same representative participant as in A. Each point represents a voxel that is well predicted by at least one language’s semantic model 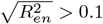 or 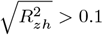. Concreteness tuning is strongly correlated between English and Chinese (*r* = .83, *p <* .001). The orthogonal regression between concreteness tuning values in English and Chinese (dotted black line) has a slope of 1.22, indicating a larger magnitude in tuning in Chinese than in English. While the cortical organization of concreteness tuning is consistent across languages, the magnitude of tuning appears to be greater in Chinese than in English. **(C)** Concreteness tuning for English (top) and Chinese (bottom) for five other participants on their cortical surface. The format is the same as in (A). The cortical organization of concreteness tuning is consistent across participants and languages.

Our results also hold when mBERT is used as the word embedding space (Fig. S11, S12). These results suggest that the cortical organization of concreteness tuning is consistent across English and Chinese.

Reading has previously been shown to be faster and more accurate for concrete words than for abstract words, a pattern described as the concreteness effect (Jessen et al. 2000). Given prior evidence that the concreteness effect is stronger in Chinese than in English (Shibahara et al. 2003), we hypothesized that the magnitude of concreteness tuning would be greater in Chinese. The magnitude of tuning can differ, even if the cortical organization of concreteness tuning is consistent between languages. Among voxels tuned towards concrete concepts in both languages (red points in Fig. 2-B), 62% of voxels have a stronger tuning towards concrete concepts in Chinese than in English. Conversely, among voxels tuned towards abstract concepts in both languages (blue points in Fig. 2-B), 54% of voxels have a stronger preference for abstract concepts in Chinese. To quantify this difference in magnitude, we fit an orthogonal regression between the concreteness tuning values in English and in Chinese (dotted black line). A slope greater than 1 indicates that the tuning magnitude is greater in Chinese than in English. This is the case for 5 out of 6 participants (1.22, 1.36, 1.15, 1.17, 0.91, 1.10 for P1-6) when fastText is used as the word embedding space (Fig. S4) and for all participants when mBERT is used (1.75, 1.76, 1.43, 1.55, 1.26, 1.32 for P1-6, Fig. S9). Thus, the magnitude of concreteness tuning is greater in Chinese than in English. That is, when a voxel is tuned towards concrete concepts, the concepts represented in Chinese have higher concreteness ratings than those represented in English. Conversely, for voxels tuned towards abstract concepts, the concepts represented in Chinese have lower concreteness ratings than those represented in English. This suggests that both the regions tuned towards concrete concepts and the regions tuned towards abstract concepts are more distinct in Chinese than in English. This may explain the observation that the concreteness effect is stronger in Chinese than in English.

### 2.3 Cross-language similarity is high for both concrete and abstract concepts

Psycholinguistic models of bilingualism predict that concrete concepts should be more similarly represented across languages than abstract concepts (Van Hell and De Groot 1998). To test this prediction, we computed the cross-language similarity of semantic representations for all voxels and compared it to the concreteness tuning. Cross-language similarity was defined as the Pearson correlation of English and Chinese estimated semantic model weights. Fig. 3-A shows cross-language similarity for one representative participant. The color of each voxel reflects the cross-language similarity. Darker colors indicate low similarity and brighter colors indicate high similarity. Voxels that are poorly predicted by both languages’ semantic models are shown in gray (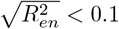 and 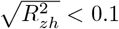). Cross-language similarity is high throughout the cortical surface and bilaterally, including in parts of the dorsolateral prefrontal cortex, Broca’s area, the high-level auditory cortex, the superior temporal sulcus, and in the lateral and medial parietal cortices. These areas correspond to both regions tuned towards concrete concepts and regions tuned towards abstract concepts (Fig. 2-A). A similar cortical distribution of cross-language similarity is observed for all other participants (Fig. S5) and when mBERT is used as the word embedding space (Fig. S13). This suggests that cross-language similarity of semantic representations is high both in regions tuned towards concrete concepts and in regions tuned towards abstract concepts.

**Fig. 3.**
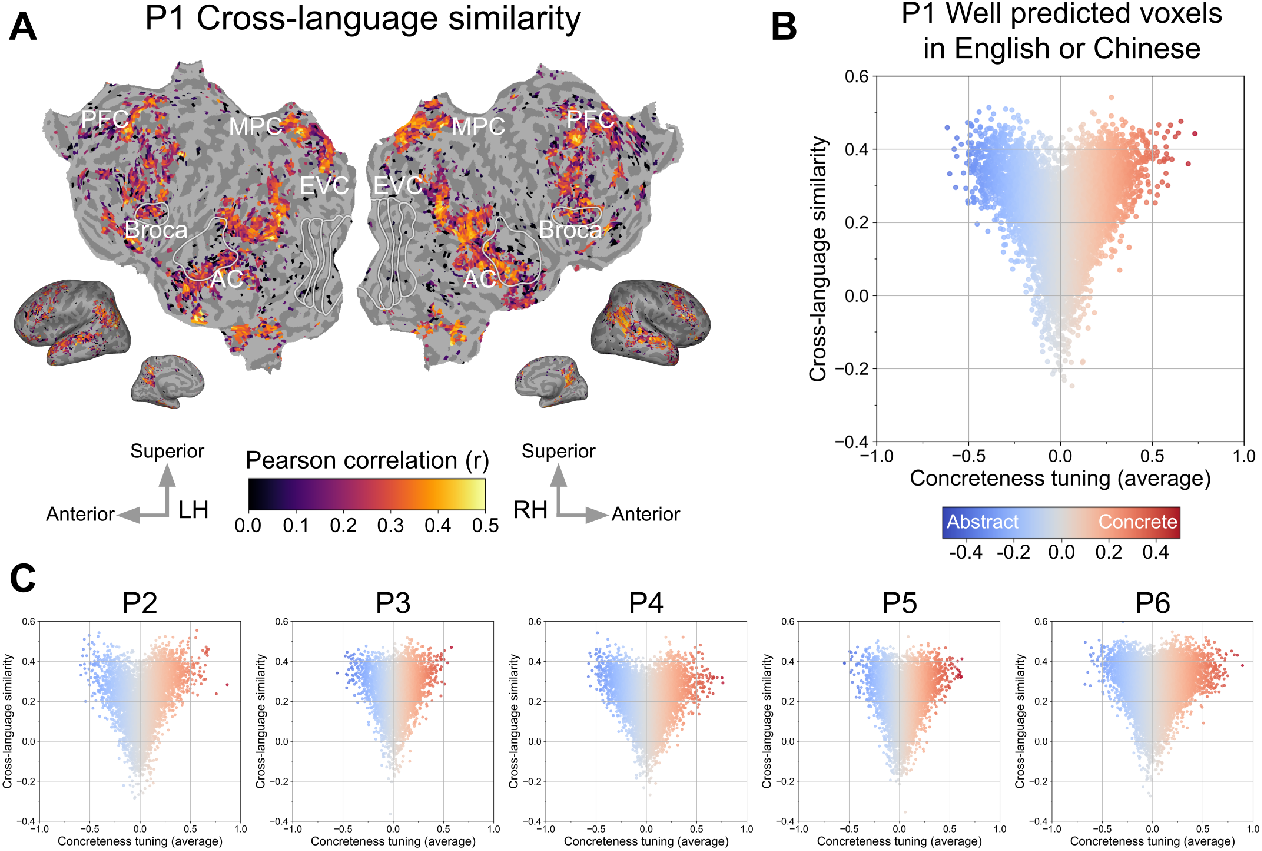
Comparison between cross-language similarity and concreteness tuning. Cross-language similarity was measured as the Pearson correlation of estimated semantic model weights between English and Chinese. **(A)** Cross-language similarity between English and Chinese semantic representations is shown for the same representative participant (P1) as in Fig. <u>2</u> on their cortical surface. Voxel color indicates the value of the Pearson correlation coefficient ranging from low similarity (dark) to high similarity (bright). Voxels with a low prediction accuracy in both languages’ semantic model are shown in gray (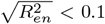 and 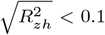). Portions of the bilateral dorsal prefrontal cortex, lateral and medial parietal cortices, high-level AC, STS and Broca’s area have high cross-language similarity. These areas coincide with both areas tuned towards abstract concepts and areas tuned towards concrete concepts in Fig. <u>2</u>-A. **(B)** Comparison of concreteness tuning (x-axis) and cross-language similarity (y-axis) in a scatter plot for the same representative participant as in (A). Each point represents a voxel that is well predicted in at least one language (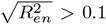 or 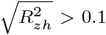). Voxels with neutral concreteness tuning (around 0) have low cross-language similarity. The more concrete (positive) or the more abstract (negative) the tuning of a voxel, the higher its cross-language similarity. Both very concrete and very abstract concepts appear to be similarly represented across languages. **(C)** Comparison of concreteness tuning and cross-language similarity for five other participants. The format is the same as in (B). Across all participants, cross-language similarity is high for both very concrete and very abstract concepts.

To quantify how cross-language similarity varies with concreteness tuning, we compared their voxelwise values. For this analysis, we took the average of the English concreteness tuning and the Chinese concreteness tuning since they are strongly correlated with each other (Fig. 2-B and Fig. S4). Fig. 3-B compares the average concreteness tuning (x-axis) and cross-language similarity (y-axis) for the same representative participant as in Fig. 2-A. Each point reflects one voxel that is well predicted in at least one language (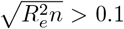 or 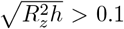). The results are consistent if we compare cross-language similarity to the English or the Chinese concreteness tuning separately (Fig. S6). Both voxels that represent highly concrete concepts (strong positive tuning) and voxels that represent highly abstract concepts (strong negative tuning) have high cross-language similarity. Voxels with a neutral concreteness tuning (around 0) have low cross-language similarity. This is observed in all participants (Fig. 3-C). We computed the correlation between concreteness tuning and cross-language similarity, separately for voxels tuned towards concrete concepts (positive concreteness tuning) and voxels tuned towards abstract concepts (negative concreteness tuning). For all participants, there is a positive correlation in concrete-tuned voxels (*r* = .53, .51, .43, .45, .38, .37, *p* < .001 by a two-sided t-test for P1-6) and a negative correlation in abstract-tuned voxels (*r* = −.59, −.48, −.41, −.54, −.39, −.40, *p <* .001 by a two-sided t-test for P1-6). Cross-language similarity increases with the absolute value of concreteness tuning, producing characteristic heart-shaped patterns. That is, voxels with a stronger tuning towards either concrete concepts or abstract concepts tend to represent more similar concepts across languages. This is also the case when mBERT is used as the word embedding space (*r* = .43, .58, .35, .47, .32, .46 in concrete-tuned voxels, *r* = −.48, −.47, −.36, −.54, −.23, .38 in abstract-tuned voxels, *p <* .001 by a two-sided t-test for P1-6, Fig. S14). This result extends previous behavioral evidence suggesting that humans process concrete words more similarly across languages than abstract words.

### 2.4 Emotional content may drive cross-language similarity of abstract concepts

We showed that cross-language similarity is high in regions tuned towards concrete concepts and in regions tuned towards abstract concepts (Fig. 3-B). This result contrasts with psycholinguistic models of bilingualism which predict lower cross-language similarity for abstract concepts (Van Hell and De Groot 1998). Based on prior evidence that abstract concept representations are driven by emotionality (Vigliocco et al. 2014), we further hypothesized that emotionality may drive cross-language similarity for abstract concepts. To test this hypothesis, we identified the emotionality dimension as a vector in the word embedding space spanning from concepts with low emotional content to concepts with high emotional content. We first identified words with low emotionality ratings and words with high emotionality ratings (Grühn 2016). We then computed the emotionality dimension as the difference between the mean embedding of the emotional words and the mean embedding of the non-emotional words. We validated the emotionality dimension by comparing the human emotionality rating of each stimulus word with the projection of the word’s embedding onto the emotionality dimension. We found that ratings and projections are strongly positively correlated (*r* = .74, *p <* .001 by a two-sided permutation test, Fig. S2), confirming that the emotionality dimension captures human judgements of emotionality. The correlation is moderately positive when using mBERT (*r* = .55, *p* < .001 by a two-sided permutation test, Fig. S10).

We then measured the emotionality tuning of each voxel by projecting the estimated semantic model weights onto the emotionality dimension. We compared emotionality tuning and cross-language similarity for abstract and concrete concepts separately. Fig. 4-A shows the comparison between emotionality tuning (x-axis) and cross-language similarity (y-axis). Results are shown separately for voxels tuned towards abstract concepts (negative concreteness tuning, upper panel) and for voxels tuned towards concrete concepts (positive concreteness tuning, lower panel) for the same representative participant as in Fig. 2 and Fig. 3. Each point represents a voxel that is well predicted in at least one language (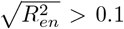 or 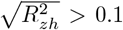). The dotted line shows the Pearson correlation between emotionality tuning and cross-language similarity. In abstract-tuned voxels, there is a moderate positive correlation between emotionality tuning and cross-language similarity (*r* = .44, *p* < .001 by a two-sided t-test). In concrete-tuned voxels, there is a weak negative correlation between emotionality tuning and cross-language similarity (*r* = −.25, *p* < .001 by a two-sided t-test). Across participants, the correlation is consistently greater in absolute value for abstract-tuned voxels (0.46, 0.50, 0.56, 0.45, 0.36 for P2-P6, Fig. 4-C) than for concrete-tuned voxels (*r* = −0.34, −0.17, −0.09, −0.16, −0.09 for P2-P6, Fig. 4-C). This difference is significant by a one-sided permutation test (*p* < .001). This is also the case when using mBERT as embedding space (for concrete-tuned voxels *r* = .57, .62, .66, .61, .37, .54 for P1-P6, for abstract-tuned voxels, *r* = −.28, −.30, −.21, −.19, −.06, −.23 for P1-P6, *p* < .001 by a two-sided t-test, Fig. S15). In other words, the more emotional an abstract concept, the more similar its representations across languages. This suggests that emotional content may drive cross-language similarity for abstract concepts.

**Fig. 4.**
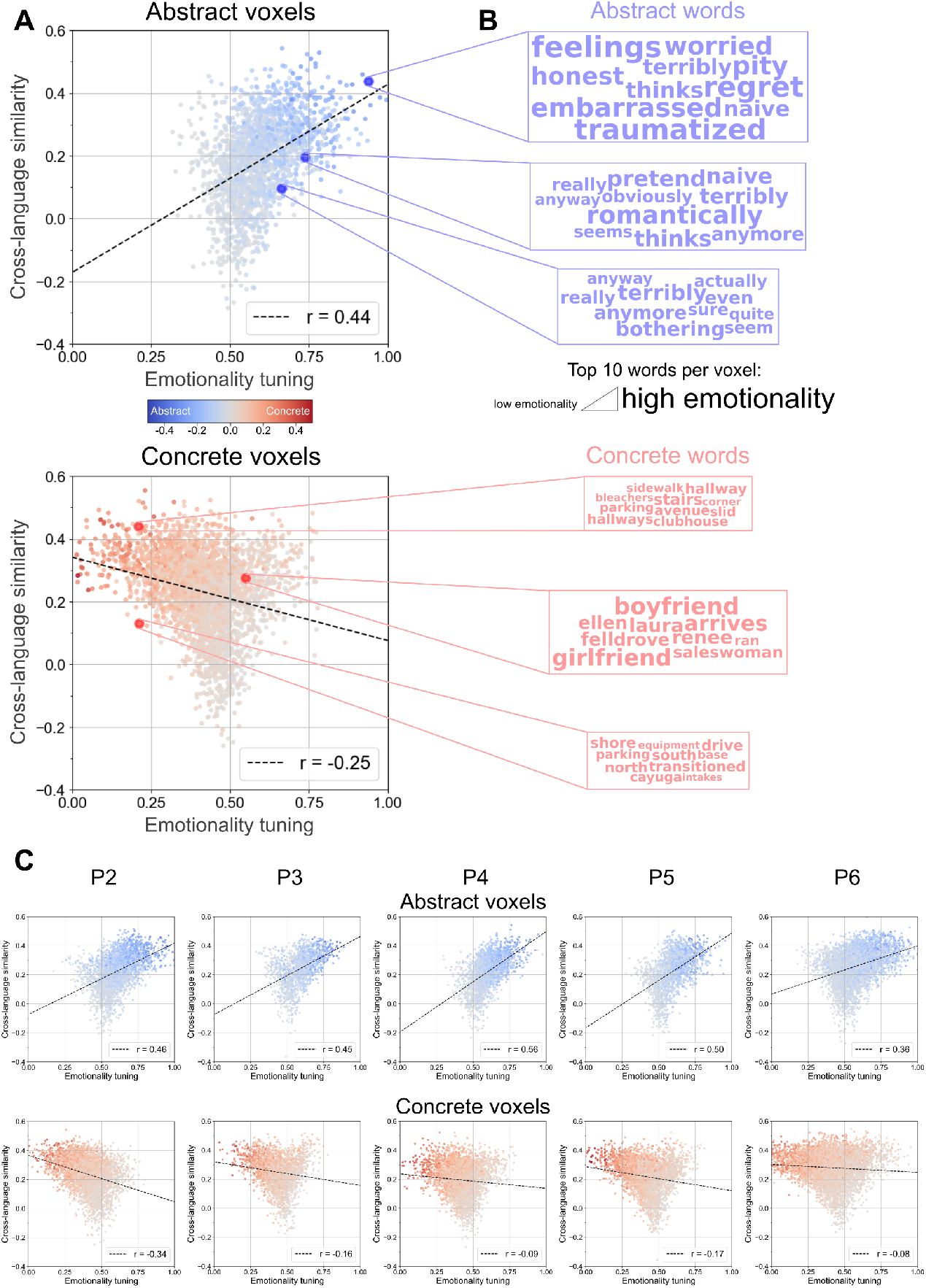
Emotionality tuning and cross-language similarity. Emotionality tuning was measured by projecting the estimated semantic model weights of each voxel onto the emotionality dimension–a vector in the embedding space spanning from concepts with low emotional content to concepts with high emotional content. Additionally, the words that elicit the highest BOLD response were identified in selected voxels. **(A)** Comparison between emotionality tuning (x-axis) and cross-language similarity (y-axis) for the same representative participant (P1) as in Fig. 2 and Fig. 3. Voxels tuned towards abstract concepts (negative concreteness tuning) are shown on the top and voxels tuned towards concrete concepts (positive concreteness tuning) are shown on the bottom. Each point represents a voxel that is well predicted in at least one language (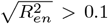 or 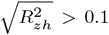). The Pearson correlation between emotionality tuning and cross-language similarity (dotted line) is stronger for abstract-tuned voxels (*r* = 0.44) than for concrete-tuned voxels (*r* = −0.25). Emotional content may drive cross-language similarity for abstract concepts. **(B)** The 10 words that elicit the highest BOLD response were selected for 3 voxels with equivalent tuning towards abstract concepts (top) and 3 voxels with equivalent tuning towards concrete concepts (bottom) but different levels of cross-language similarity. The size of each word is proportional to its emotionality tuning. For concrete words, there is no pattern across levels of cross-language similarity. For abstract words, the higher the cross-language similarity of a voxel, the more emotional the words it represents. This illustrates the result that emotional content may drive cross-language similarity for abstract concepts. **(C)** Comparison between emotionality tuning and cross-language similarity for five other participants. The format is the same as in (A). Across all participants, emotional content may drive cross-language similarity for abstract concepts.

To illustrate the correlation between emotionality tuning and cross-language similarity, we identified the words that elicit the highest blood oxygen level dependent (BOLD) response in selected voxels. For both abstract-tuned and concrete-tuned voxels, we chose three voxels with equivalent concreteness tuning but varying levels of cross-language similarity. Fig. 4-B shows the ten closest words for each voxel. The size of each word is proportional to its emotionality tuning. For voxels tuned towards concrete concepts, there is no strong correspondence between cross-language similarity and emotionality tuning. In contrast, for voxels tuned towards abstract concepts, the higher the cross-language similarity of a voxel, the more emotional the word it represents. For example, the voxel with highest cross-language similarity has almost exclusively extremely emotional words such as “feelings”, “regret” or “traumatized”. This is consistent with the observation that, in voxels tuned towards abstract concepts, emotional content may drive similarity of representations across languages.

Additionally, a direct comparison of concreteness tuning and emotionality tuning across all voxels shows that abstract-tuned voxels have a higher emotional content (Fig. S8, Fig. S17 with mBERT). This is also shown in Fig. 4-B, where all concrete words except for a few exceptions (“girlfriend” or “boyfriend”) have much lower emotionality tuning than abstract words. Overall, abstract-tuned voxels have a stronger emotionality tuning, and this emotionality tuning may drive cross-language similarity. This suggests that emotional content plays a crucial role in the representation of abstract concepts across languages.

## 3 Discussion

This study investigated how concreteness influences the similarity of semantic representations across languages in bilingual brains. We used voxelwise encoding models to estimate the semantic information represented in each voxel, in English and in Chinese separately (Fig. 1-A). First, we measured the tuning of each voxel towards concrete and abstract concepts in each language (Fig. 1-B). We found that the cortical organization of concreteness tuning is consistent across languages (Fig. 2). Second, we measured the similarity of the semantic representations across languages in each voxel (Fig. 1-C). Cross-language similarity is high both in voxels tuned towards concrete concepts and voxels tuned towards abstract concepts (Fig. 3). Last, we evaluated the emotional content of semantic representations in each voxel. Abstract-tuned voxels that encode similar concepts across languages are more likely to represent emotional words (Fig. 4).

A large body of work primarily using the English language has studied the cortical organization of concreteness tuning (see (Wang et al. 2010) for a meta-analysis). These studies consistently show that concrete and abstract concepts are represented in distinct regions. In line with grounded cognition theories (Barsalou 2008), representations of concrete concepts are located closer to perceptual systems, whereas abstract concepts are primarily represented in areas typically associated with language. Cortical regions that represent a broad range of concepts, such as the dorsolateral prefrontal cortex, are involved in representations of both concrete and abstract concepts (Binder et al. 2005; Vigliocco et al. 2014). Our results are consistent with these findings. We find that posterior areas in the lateral and medial parietal cortices are tuned towards concrete concepts (Fig. 2-A). These posterior regions lie at the edge of object-selective areas immediately anterior to the early visual cortex. In these areas, visual and linguistic semantic information converge (Popham et al. 2021), supporting the representation of concrete concepts grounded in perceptual experience. In contrast, we find that abstract concepts are represented mainly in Broca’s area, the high-auditory cortex and the superior temporal sulcus (Fig. 2-A). These areas are typically associated with language (Fedorenko et al. 2016; Caucheteux et al. 2023). We find that the dorsolateral prefrontal cortex is tuned toward both concrete and abstract concepts. These results are consistent with dual coding theory (Paivio 1991) which suggests that abstract concepts are represented only in verbal memory whereas concrete concepts also engage visual memory.

Our study shows that the cortical organization of concrete and abstract concepts is not unique to English and is consistent with Chinese (Fig. 2). Regions tuned towards concrete concepts in one language are also tuned towards concrete concepts in the other, and the same holds for abstract concepts. This supports prior fMRI studies showing commonalities in semantic representation across languages (Buchweitz et al. 2012; Yang et al. 2017b,a; Dehghani et al. 2017; Chen et al. 2024a) and suggests that concreteness is a semantic category that is organized similarly across languages in the bilingual brain.

Prior work has shown that the cortical representations of some semantic categories can be differently modulated in each language (Chen et al. 2024a). We observe a similar pattern in our results: while the cortical organization of concreteness tuning is consistent across languages, its magnitude is larger in Chinese than in English (Fig. 2-B). This suggests that representations of concrete and abstract concepts are more distinct in Chinese than in English. There are two potential reasons for this result. First, Chinese uses a logographic writing system whereas English uses an alphabetic writing system. Therefore, the difference in concreteness tuning could reflect differences in brain representations of different writing systems. Indeed, the effect of concreteness on behavioral tasks such as word naming or on cortical representations has been shown to be stronger when reading logographs than alphabets (Shibahara et al. 2003; Chee et al. 2000). Second, all participants were native Chinese speakers who reached proficiency in English. Thus, the greater magnitude in Chinese concreteness tuning could reflect differences in native versus non-native language comprehension. Our analysis methodology is language agnostic, and in the future we hope to work with a broader range of bilingual participants or other language pairs to determine how our findings extend to other language backgrounds and writing systems.

Our findings have implications for psycholinguistic models of bilingualism, where concreteness is an important factor (Pavlenko 2009). For example, based on behavioral data showing that concrete words are faster to translate (Basnight-Brown and Altarriba 2016; Tokowicz and Kroll 2007), the distributed conceptual feature model predicts that concrete concepts are more similarly represented across languages than abstract concepts (de Groot 1992). Higher similarities for concrete concepts than for abstract concepts have also been observed in semantic representations in the brain, but only across monolingual speakers of the same language (Botch and Finn 2024) and not in bilingual brains. Our results partly diverge from these findings. We show that both regions tuned towards concrete concepts and regions tuned towards abstract concepts have high cross-language similarity (Fig. 3-B). In fact, cross-language similarity increases as voxels represent concepts that are either more concrete or more abstract.

At least two factors could explain the divergence between our results and the distributed conceptual feature model. One stems from the type of experiment, and one from the underlying concepts. First, we use naturalistic stimuli, while psycholinguistic experiments often use controlled stimuli where words appear isolated or in short sentences with no broad context. Prior work has shown that brain representations of concrete and abstract concepts depend on the context in which they appear (Kewenig et al. 2023). They suggest that when abstract concepts appear in the context of objects, they are represented more in regions that usually process concrete concepts. Conversely, concrete concepts appearing out of context are represented more in regions usually tuned towards abstract concepts. This could explain why we show higher cross-language similarity for abstract concepts than in experiments with controlled stimuli, as the abstract concepts in our stimuli appear within a natural context.

A second factor is the type of abstract concepts for which we find high cross-language similarity. Our results show that abstract concepts with higher cross-language similarity tend to have greater emotional content (Fig. 4). In line with previous findings, we also show that abstract concepts are more emotional than concrete concepts overall (Fig. S8) (Kousta et al. 2011). This indicates that emotional content plays a particularly important role in the representation of abstract concepts. Across mono-lingual speakers of English and Chinese, prior work has found that representations of abstract concepts are similar along certain dimensions related to the self or social content (Vargas and Just 2022). Within language, it has been shown that representations of abstract concepts are grounded in affective experience through emotions (Vigliocco et al. 2014). Our results suggest that emotions also ground concepts across languages. Some researchers have proposed considering concrete, abstract and emotion words as three separate categories as they lead to different behavioral results in bilingual representations (Altarriba 2003; Pavlenko 2008). Future work could study the emotionality dimension orthogonally to concreteness to isolate their individual contributions.

In this study, we investigated how concreteness influences the similarity of semantic representations across languages in bilingual brains. We found that the organization of tuning towards concrete and abstract concepts is consistent across English and Chinese. As predicted by psycholinguistic models of bilingualism, we observed that cross-language similarity is high for concrete concepts. In addition, we showed that cross-language similarity is also high for abstract concepts, particularly for those with emotional content. While sensory experience grounds concrete concepts across languages, emotional experience may do the same for abstract concepts.

## 4 Methods

### 4.1 Participants

Brain responses were recorded from six bilingual participants who are fluent in both Mandarin Chinese (L1; native) and English (L2; non-native). Participants began learning English between the ages of 2 and 11. At the time of the experiment they had been living and studying or working in an English-speaking country for between 5 to 12 years and were using both languages on a daily basis. Language proficiency was evaluated with the Language Experience and Proficiency Questionnaire (LEAP-Q) (Kaushanskaya et al. 2020) and the Language History Questionnaire (LHQ3) (Li et al. 2020). All participants were healthy and had normal or corrected-to-normal vision. All participants were right handed or ambidextrous according to the Edinburgh handedness inventory (Oldfield 1971). The data were originally collected for separate experiments; we refer the readers to (Chen et al. 2024a) for additional information on the participants’ use of each language.

### 4.2 Stimuli

Eleven narratives from The Moth Radio Hour were used as stimuli. Each narrative lasts between 10 to 15 minutes and consists of a speaker telling an autobiographical story in English. The audio recordings were manually transcribed and the written transcriptions were aligned to match the audio versions. The English narratives were translated into Chinese by a professional translator. They were read by a professional actor to obtain natural speech rates and then transcribed and aligned to these audio recordings. Simplified Chinese characters were used for stimulus presentation. Further information about the stimulus presentation format can be found in (Chen et al. 2024a). Ten narratives were used to estimate the encoding models and the eleventh held-out narrative was used to evaluate the estimated models.

### 4.3 fMRI data collection and preprocessing

Whole-brain MRI data were collected on a 3T siemens TIM trio scanner at the UC Berkeley Brain Imaging Center. A 32-channel Siemens volume coil was used. Functional scans were collected using a T2*-weighted gradient-echo EPI with repetition time (TR) 2.0045s, echo time (TE) 35ms, flip angle 74°, voxel size 2.24×2.24×4.1 mm (slice thickness 3.5mm with 18% slice gap), matrix size 100×100, and field of view 224×224 mm. Thirty axial slices were prescribed to cover the entire cortex and were scanned in interleaved order. A custom-modified bipolar water excitation radiofrequency (RF) pulse was used to prevent contamination from fat signals. Anatomical data were collected using a T1-weighted multi-echo MP-RAGE sequence on the same 3T scanner. To minimize head motion during scanning, participants wore a personalized head case that precisely fit the shape of their head (Gao 2015; Power et al. 2019). Preprocessing of the data included motion correction using FLIRT from FSL (Jenkinson et al. 2002) and removal of noise from motion, respiratory and cardiac signals with CompCor (Behzadi et al. 2007). Further specifics about preprocessing can be found in (Chen et al. 2024a). Responses were z-scored across time to unit variance centered around zero for each voxel and narrative. The first and last 10 TRs of each narratives were discarded to take into account the silence at the beginning and end of each scan and the non-stationarity in brain responses.

### 4.4 Feature spaces

#### 4.4.1 Semantic features

To represent the semantic information of the stimulus words, we used vectors from a word embedding space. We selected lexical vectors from fastText (Bojanowski et al. 2017). We used the version of fastText that is aligned across languages to facilitate comparisons between English and Chinese (Joulin et al. 2018). These vectors map each word to a 300-dimensional semantic space. We additionally validated our results with a separate word embedding space, multilingual BERT (mBERT) (Devlin et al. 2019). mBERT is a Transformer-based language model with 12 layers and hidden layer size of 768. We obtained a contextual vector for each word by providing the current word and 10 previous words as context to mBERT. We used a context size of 10 as it was shown to produce accurate predictions of brain responses (Toneva and Wehbe 2019; Negi Name et al. 2025). We used layer 0 because the way we define the concreteness and emotionality dimensions uses words in a lexical manner, and later layers are more contextualized. fastText and mBERT have been shown to capture information about word concreteness (Charbonnier and Wartena 2019; Wartena 2024).

#### 4.4.2 Low-level sensory features

To account for the effects of sensory information on brain responses, we created seven feature spaces that reflect low-level characteristics of the stimulus. These features represent the word count, letter count for English or character count for Chinese, single phonemes, diphones, triphones (Gong et al. 2023), visual and spatial motion features (motion energy) (Adelson and Bergen 1985; Watson and Ahumada Jr 1985; Nishimoto and Gallant 2011) and intermediate features that capture orthographic similarities by measuring pixel overlap between words (Gong 2024).

#### 4.4.3 Feature space preprocessing

To align the stimulus feature spaces with the fMRI temporal resolution, we processed each feature space using a three-lobe Lanczos low-pass filter with a 0.25 Hz cut-off frequency. We then accounted for the temporal delay between neural activity and the measured BOLD response by implementing a finite impulse response (FIR) model. The FIR model created time-shifted versions of each feature space at delays of 2, 4, 6, and 8 seconds to capture the hemodynamic response of individual voxels, following prior work (Huth et al. 2016; Deniz et al. 2019; Nakai et al. 2021).

### 4.5 Voxelwise encoding model fitting

To determine which features are represented in each voxel, we estimate voxelwise encoding models (Naselaris et al. 2011; Huth et al. 2016; De Heer et al. 2017; Deniz et al. 2019; Popham et al. 2021; Lamarre et al. 2022; Deniz et al. 2023; Chen et al. 2024b,Chen et al. 2024a; Visconti di Oleggio Castello et al. 202; Negi Name et al. 2025). Each model is defined by a set of regression weights, which characterize the BOLD responses of a single voxel as a linear combination of features from a given feature space. We used banded ridge regression to estimate the regression weights (Nunez-Elizalde et al. 2019). Banded ridge regression is a variant of standard ridge regression that allows different regularization parameters for each feature space, and therefore can avoid biases caused by differences in feature distributions. Mathematically, banded ridge regression estimates the weights for each feature space by solving: 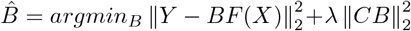 where *B* is the mapping of dimension (dimension 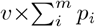), *F* ^*′*^(*X*) is the concatenated feature of *m* delayed feature spaces (each of dimension *p*_*i*_), *Y* is the brain responses (dimension *v* × *n*) and *C* the diagonal matrix of regularization parameters, with *v* the number of voxels and *n* the number of TRs. We used 5-fold cross-validation to optimize the *C* for each feature space and each voxel using the Himalaya Python package (La Tour et al. 2022). We used a random search procedure with 1000 normalized hyper-parameters candidates randomly sampled from a dirichlet distribution and scaled by 21 log-spaced values ranging from 10^*−*^10 to 10^10^. We selected the hyperparameters that minimize the squared error.

#### 4.5.1 Stepwise regression procedure

To control for correlations between semantic and low-level sensory features in the stimulus, we used a stepwise regression procedure. We first jointly estimated encoding models that predict BOLD responses from seven low-level sensory features (see 4.4.2). We used only data from the train narratives to estimate the low-level models. We then predicted BOLD responses to the train and test narratives. We subtracted the predicted BOLD responses 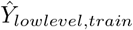 and 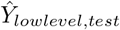 from the recorded BOLD responses *Y*_*train*_, *Y*_*test*_. We z-scored the residual BOLD responses 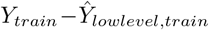 and 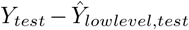 to unit variance centered around zero for each voxel. We used the z-scored residual BOLD responses to estimate encoding models with the semantic features.

### 4.6 Projections in the word embedding space

#### 4.6.1 Concreteness

To measure the preference of each voxel for concrete or abstract concepts (concreteness tuning), we linearly projected each voxel’s estimated semantic model weights onto the concreteness dimension. As the dimensions of the semantic spaces of fastText or multilingual BERT do not have an explicit meaning, this concreteness dimension was identified as in (Grand et al. 2022). In English and in Chinese, we selected a list of words which denote highly concrete concepts and a list of words which denote highly abstract concepts based on databases of human ratings of word concreteness (Brysbaert et al. 2014; Xu and Li 2020). For each language, we computed the mean of the semantic vectors of the concrete words and the mean of the semantic vectors of the abstract words, then defined the concreteness dimension as the vector difference (concrete minus abstract). Any vector in the word embedding space (word vector or estimated semantic model weights) can be projected onto this concreteness dimension to estimate the concreteness of the corresponding concept. These dimensions were validated by comparing the human ratings of concreteness with the projections (Fig. S1, Fig. S6). We computed the Pearson correlation between human rating and projection for all the stimulus words and found the highest correlations when using the 500 most concrete and 500 most abstract words from the databases to construct the concreteness dimension. We therefore used these words to construct the concreteness dimension in all analyses. As the projections are unitless, we scaled them to have values approximately between −1 and 1.

#### 4.6.2 Human ratings extension

The coverage of the Chinese concreteness rating database (Xu and Li 2020) on our dataset was very low as it only covers 2-character words. To extend it for the validation of the concreteness dimension step (comparison of human ratings of concreteness with projections), we predicted ratings for the remaining words using BERT embeddings as in (Turton et al. 2021).

#### 4.6.3 Emotionality

To evaluate the emotional content of the concepts represented in each voxel (emotionality tuning), we linearly projected each voxel’s estimated semantic model weights onto the emotionality dimension. This dimension was identified using the same technique we used for the concreteness dimension (see 4.6.1 above). We selected words which denote concepts with high emotionality and concepts with low emotionality based on a database of human ratings of word emotionality in English (Grühn 2016). This dimension was validated by comparing the human ratings of emotionality with the projections (Fig. S2, Fig. S7). We computed the Pearson correlation between human rating and projection for all the stimulus words and found the highest correlations when using top 100 and bottom 100 words from the database to construct the emotionality dimension. We therefore used these words to construct the emotionality dimension in all analyses. As the projections are unitless, we scaled them to have values approximately between −1 and 1.

### 4.7 Highest-response words

To select the words that elicit the highest-response in a given voxel, we used the voxel’s estimated semantic model weights. We measured the distance between that voxel’s semantic weights and the word vectors of all the stimulus words by computing their dot product. We selected the ten words with the smallest distance to the voxel’s semantic weights.

### 4.8 Magnitude of concreteness tuning

To measure the relationship between the magnitude of concreteness tuning in English and in Chinese, we used orthogonal regression because we do not assume that tuning in one language predicts tuning in the other. Rather, we consider that both are noisy estimates of an underlying relationship.

### 4.9 Voxel selection

To study only voxels where the estimated semantic model weights accurately describe the semantic information represented, we selected voxels with high prediction accuracy (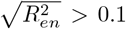 or 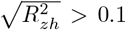). Selecting voxels based on significance (Fig. S9, Fig. S18) instead of prediction accuracy produces similar results.

## Supplementary information

This article has an accompanying supplementary information file.

## Acknowledgements

We thank Christine Tseng and Anuja Negi for valuable discussions. This work was funded by grants from the National Science Foundation (NAT-1912373) and the German Federal Ministry of Education and Research (BMBF 01GQ1906). CC was supported in part by an NSF GRFP and an IBM PhD fellowship.

