## Supplementary figures for "Concreteness shapes semantic representations in bilingual brains"

### Supplementary information

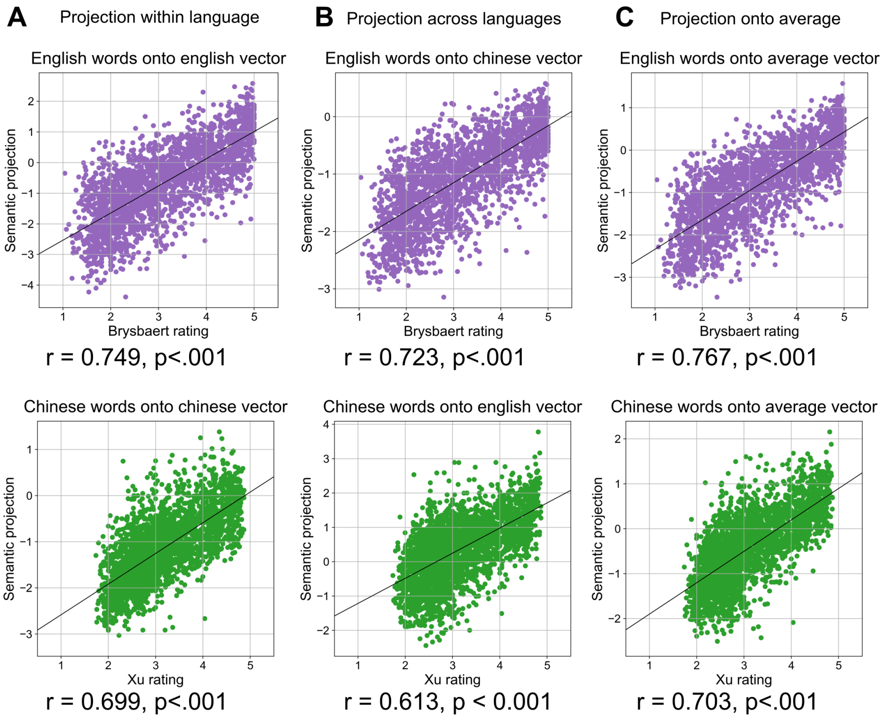


**Fig. S1 Projections of stimulus word embeddings onto the concreteness dimension in fastText space.** To validate the precision of the concreteness dimension we identified in the word embedding space, we compared the human rating of concreteness (x-axis) with the projection of the word's embedding onto the concreteness dimension (y-axis) for all stimulus words. Each point represents one word. English words are plotted in purple and Chinese words are plotted in green. **(A)** shows the projections onto the dimension of the same language, **(B)** shows the projections onto the dimension of the other language and **(C)** shows the projections onto the average of the English and Chinese dimensions. Pearson correlation (r) between human ratings and projections is high both within and across languages. This suggests that our method is a reliable way to measure the concreteness of a concept and that English and Chinese word embedding spaces are well aligned.

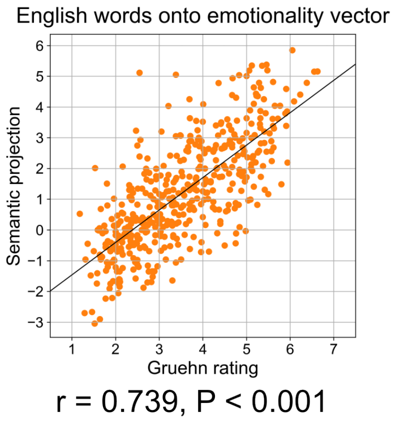


**Fig. S2 Projection of stimulus word embeddings onto the emotionality dimension in fastText space.** To validate the precision of the emotionality dimension we identified in the word embedding space, we compared the human rating of emotionality (x-axis) with the projection of the word's embedding onto the emotionality dimension (y-axis) for all stimulus words. Each point represents one word. Pearson correlation (r) between human ratings and projections is high. This suggests that our method is a reliable way to measure the emotionality of a concept.

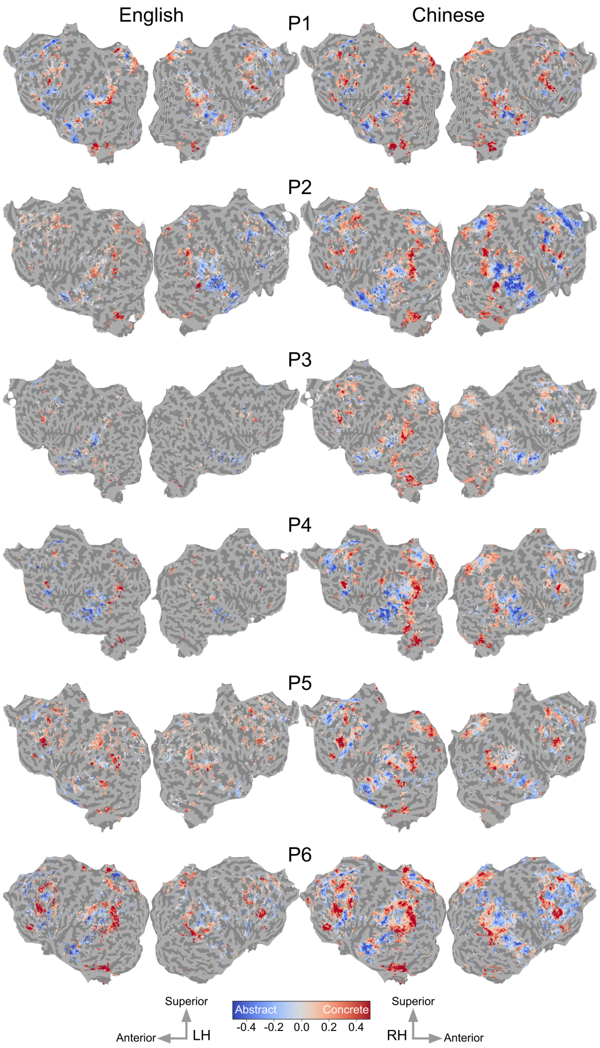


**Fig. S3 Concreteness tuning in English and Chinese for all participants in fastText space.** Concreteness tuning is shown on the flattened cortical surface of each participant. Voxel color indicates the value of concreteness tuning ranging from abstract (blue) to concrete (red). Voxels with a low prediction accuracy in the semantic model are shown in gray ($\sqrt{R^2} < 0.1$). The cortical organization of concreteness tuning is consistent across participants and languages.

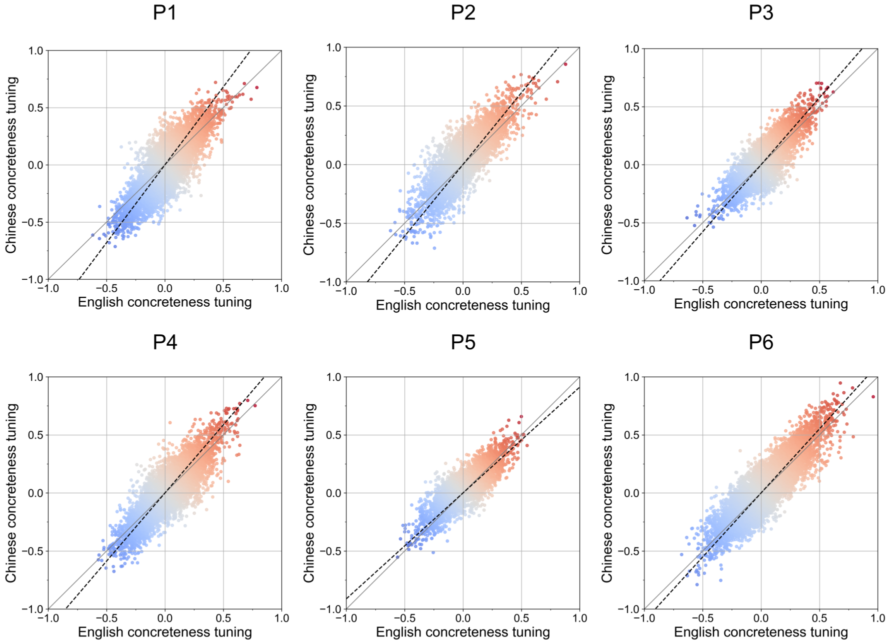


**Fig. S4 Comparison of concreteness tuning between English (x-axis) and Chinese (y-axis) for all participants in fastText space.** Each point represents a voxel that is well predicted in at least one language ($\sqrt{R_{en}}^2 > 0.1$ or $\sqrt{R_{zh}}^2 > 0.1$). Concreteness tuning is highly correlated between English and Chinese for all participants ($r = 0.83, 0.82, 0.82, 0.85, 0.83, 0.89, p < .001$ for P1-6). This suggests that the cortical organization of concreteness tuning is consistent across languages. The orthogonal regression (dotted black line) has a slope of higher than one for 5 out of 6 participants ($1.22, 1.36, 1.15, 1.17, 0.91, 1.10$ for P1-6), indicating a greater magnitude of tuning in Chinese. While the organization of concreteness tuning is consistent across languages, it has a greater magnitude in Chinese than in English.

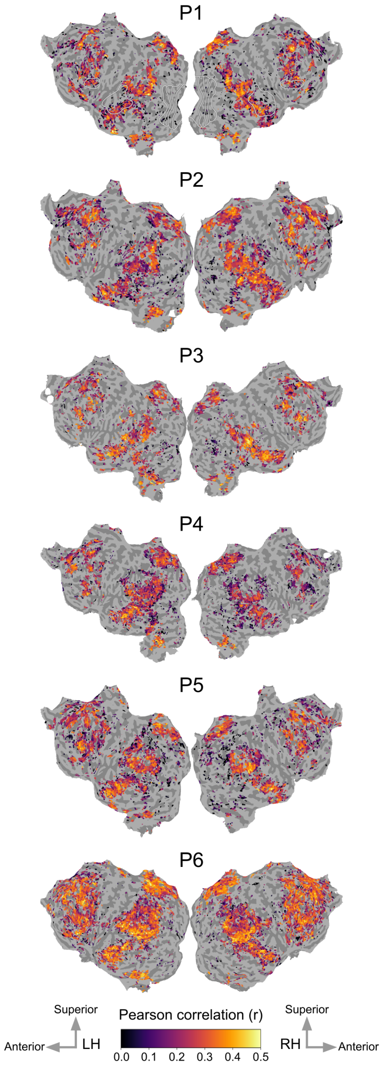


**Fig. S5 Cross-language similarity for all participants in fastText space.** Cross-language similarity between English and Chinese semantic representations is shown for all participants on their cortical surface. Voxel color indicates the value of the Pearson correlation coefficient ranging from low similarity (dark) to high similarity (bright). Voxels with a low prediction accuracy in both languages' semantic models are shown in gray ($\sqrt{R_{en}^2} < 0.1$ and $\sqrt{R_{zh}^2} < 0.1$). Portions of the dorsolateral prefrontal cortex, the lateral and medial parietal cortices, the high-level auditory cortex, the superior temporal sulcus and Broca's area have high cross-language similarity.

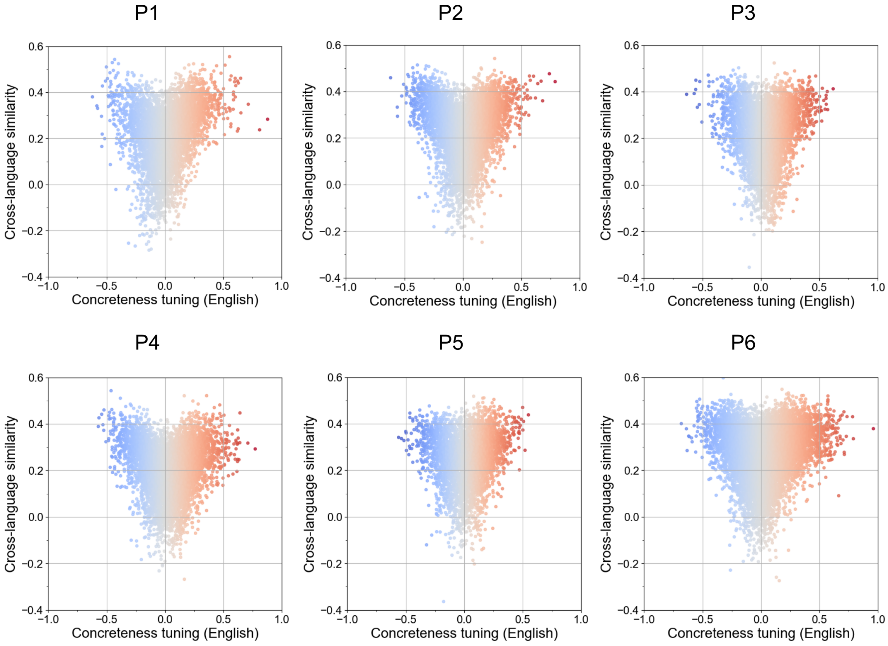


**Fig. S6 Comparison between concreteness tuning in English (x-axis) and cross-language similarity (y-axis) for all participants in fastText space.** Each point represents a voxel that is well predicted in at least one language ($\sqrt{R_{en}}^2 > 0.1$ or $\sqrt{R_{zh}}^2 > 0.1$). The results are consistent with using the average of the English and Chinese concreteness tuning (Fig. 3). This suggests that the cortical organization of concreteness tuning is consistent across languages.

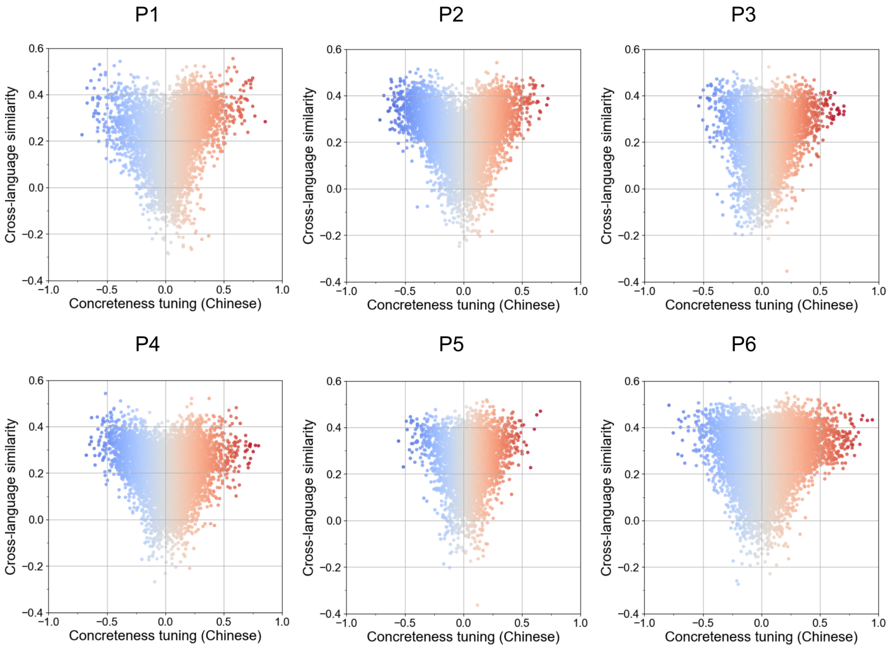


**Fig. S7 Comparison between concreteness tuning in Chinese (x-axis) and cross-language similarity (y-axis) for all participants in fastText space.** Each point represents a voxel that is well predicted in at least one language ($\sqrt{R_{en}}^2 > 0.1$ or $\sqrt{R_{zh}}^2 > 0.1$). The results are consistent with using the average of the English and Chinese concreteness tuning (Fig. 3). This suggests that the cortical organization of concreteness tuning is consistent across languages.

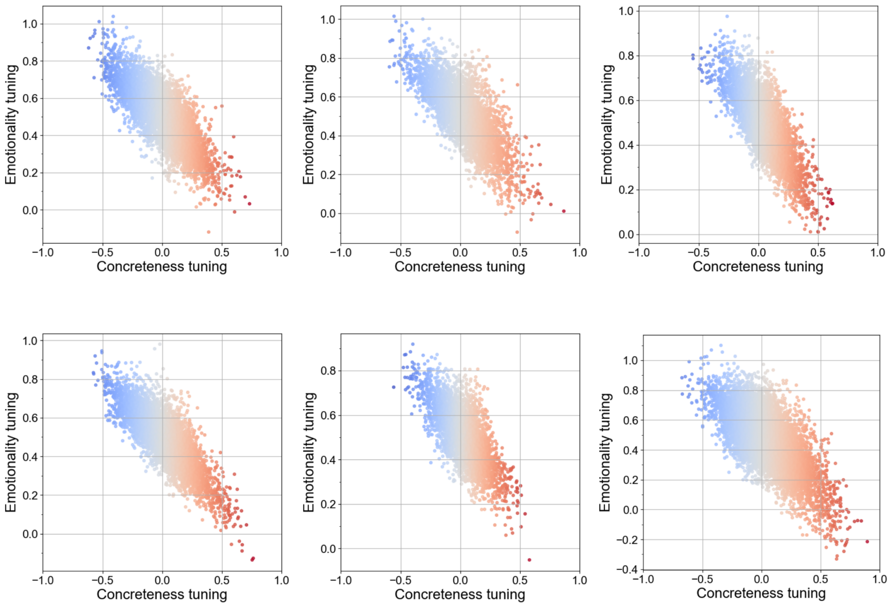


**Fig. S8 Comparison between concreteness tuning (x-axis) and emotionality tuning (y-axis) for all participants in fastText space.** Each point represents a voxel that is well predicted in at least one language ($\sqrt{R_{en}}^2 > 0.1$ or $\sqrt{R_{zh}}^2 > 0.1$). There is a negative correlation between concreteness tuning and emotionality tuning. This suggests that regions tuned towards abstract concepts represent concepts with greater emotional content.

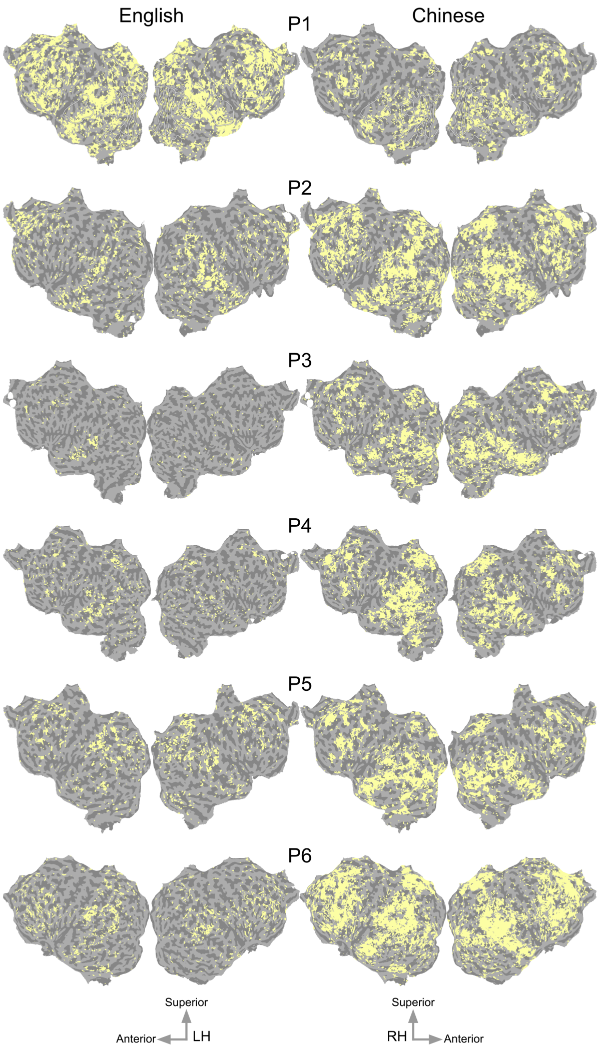


**Fig. S9 Statistical significance of prediction accuracy for each participant in each language in fastText space.** The estimated semantic model weights of the fastText embedding space were used to predict voxel responses on a held-out story. Prediction accuracy was computed as the coefficient of determination ($R^2$) between predicted and recorded brain response on the held-out story. The statistical significance of prediction accuracy of each voxel was determined by comparing the prediction accuracy of the estimated models to the prediction accuracy in predicting permuted data. Voxels that were significantly predicted are shown on the flattened cortical surface of each participant, separately for each language. Voxels shown in yellow were significantly predicted (one-sided p¡.05, FDR corrected with a Benjamini-Hochberg correction for multiple comparisons).

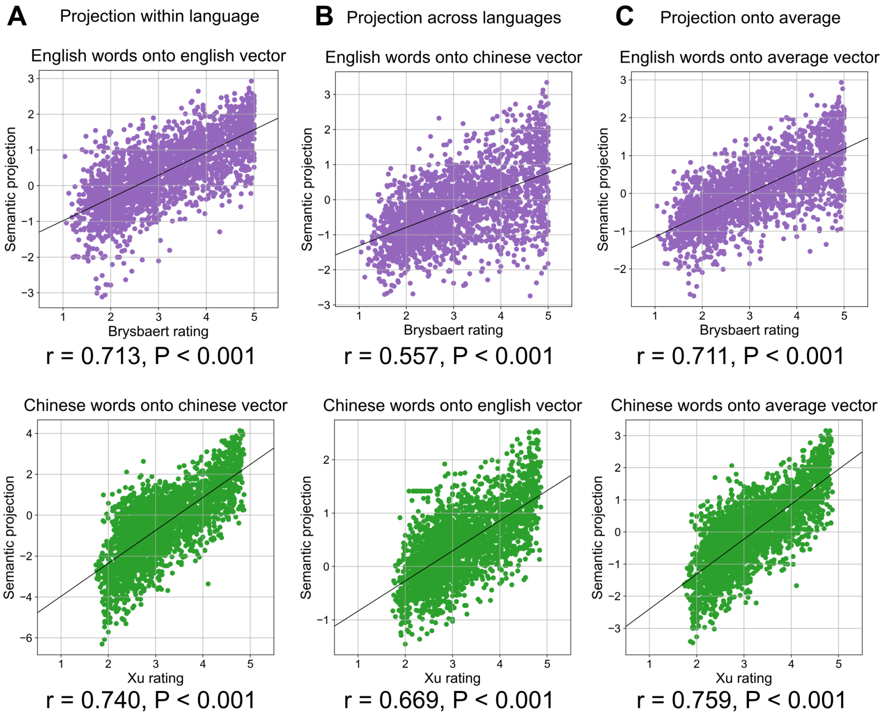


**Fig. S10 Projections of stimulus word embeddings onto the concreteness dimension in multilingual BERT.** To validate the precision of the concreteness dimension we identified in the word embedding space, we compared the human rating of concreteness (x-axis) with the projection of the word's embedding onto the concreteness dimension (y-axis) for all stimulus words. Each point represents one word. English words are plotted in purple and Chinese words are plotted in green. **(A)** shows the projections onto the dimension of the same language, **(B)** shows the projections onto the dimension of the other language and **(C)** shows the projections onto the average of the English and Chinese dimensions. Pearson correlation (r) between human ratings and projections is high both within and across languages. This suggests that our method is a reliable way to measure the concreteness of a concept and that English and Chinese word embedding spaces are well aligned.

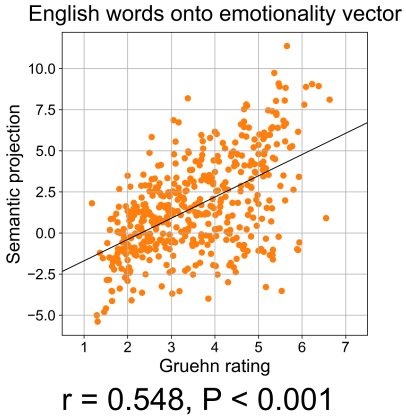


**Fig. S11 Projection of stimulus word embeddings onto the emotionality dimension in multilingual BERT.** To validate the precision of the emotionality dimension we identified in the word embedding space, we compared the human rating of emotionality (x-axis) with the projection of the word's embedding onto the emotionality dimension (y-axis) for all stimulus words. Each point represents one word. Pearson correlation (r) between human ratings and projections is high. This suggests that our method is a reliable way to measure the emotionality of a concept.

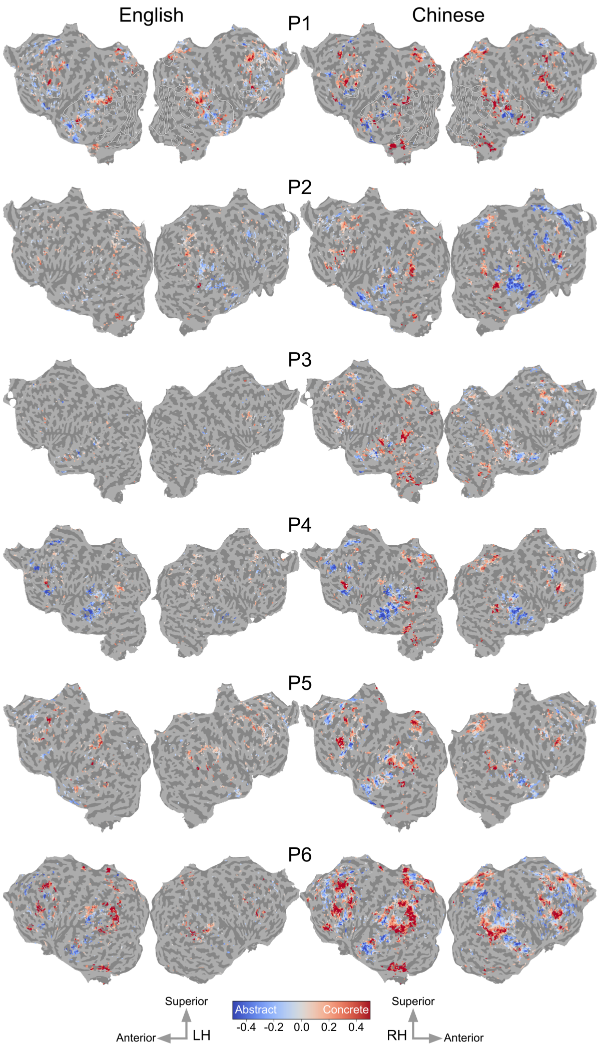


**Fig. S12 Concreteness tuning in English and Chinese for all participants in multilingual BERT.** Concreteness tuning is shown on the flattened cortical surface of each participant. Voxel color indicates the value of concreteness tuning ranging from abstract (blue) to concrete (red). Voxels with a low prediction accuracy in the semantic model are shown in gray ($\sqrt{R^2} < 0.1$). The cortical organization of concreteness tuning is consistent across participants and languages.

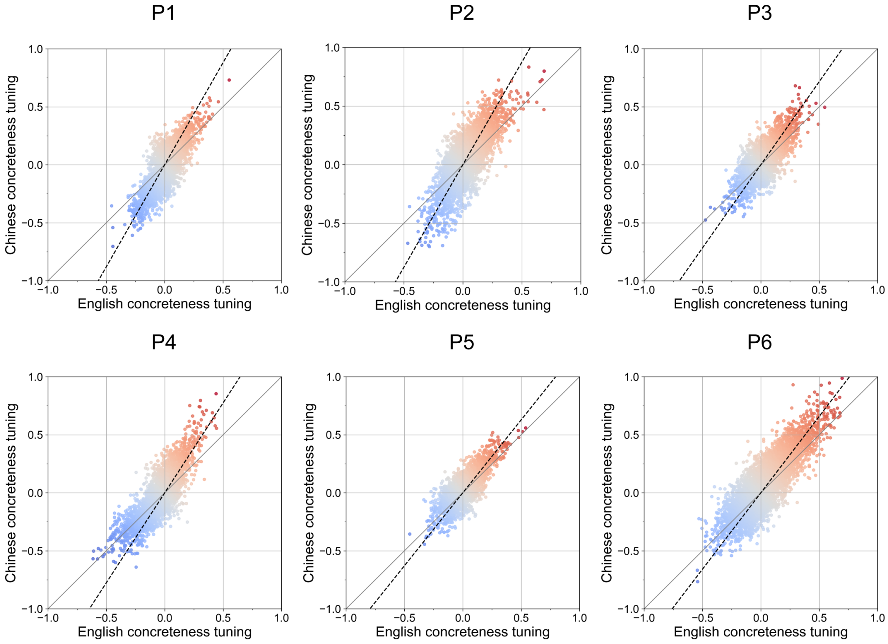


**Fig. S13 Comparison of concreteness tuning between English (x-axis) and Chinese (y-axis) for all participants in multilingual BERT.** Each point represents a voxel that is well predicted in at least one language ($\sqrt{R_{en}}^2 > 0.1$ or $\sqrt{R_{zh}}^2 > 0.1$). Concreteness tuning is highly correlated between English and Chinese for all participants ($r = 0.78, 0.80, 0.74, 0.83, 0.79, 0.84, p <$ .001 for P1-6). This suggests that the cortical organization of concreteness tuning is consistent across languages. The orthogonal regression (dotted black line) has a slope of higher than one for 5 out of 6 participants ($1.75, 1.76, 1.43, 1.55, 1.26, 1.32$ for P1-6), indicating a greater magnitude of tuning in Chinese. While the organization of concreteness tuning is consistent across languages, it has a greater magnitude in Chinese than in English.

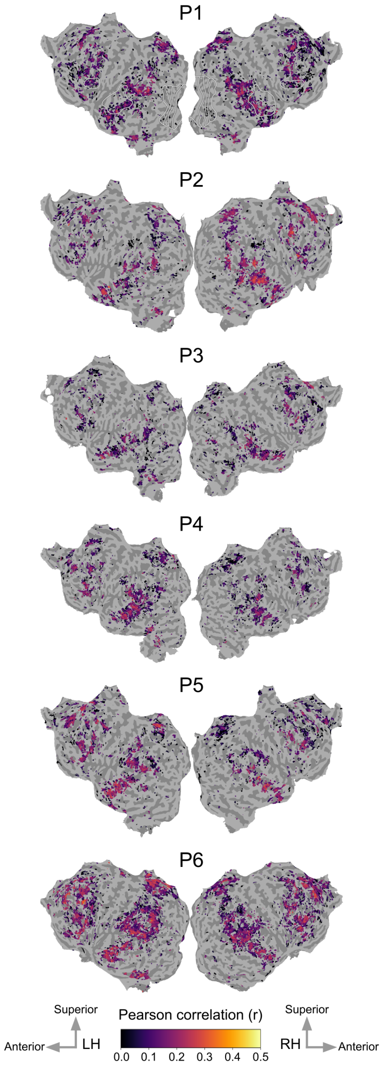


**Fig. S14 Cross-language similarity for all participants in multilingual BERT.** Cross-language similarity between English and Chinese semantic representations is shown for all participants on their cortical surface. Voxel color indicates the value of the Pearson correlation coefficient ranging from low similarity (dark) to high similarity (bright). Voxels with a low prediction accuracy in both languages' semantic models are shown in gray ($\sqrt{R_{en}^2} < 0.1$ and $\sqrt{R_{zh}^2} < 0.1$). Portions of the dorsolateral pre-frontal cortex, the lateral and medial parietal cortices, the high-level auditory cortex, the superior temporal sulcus and Broca's area have high cross-language similarity.

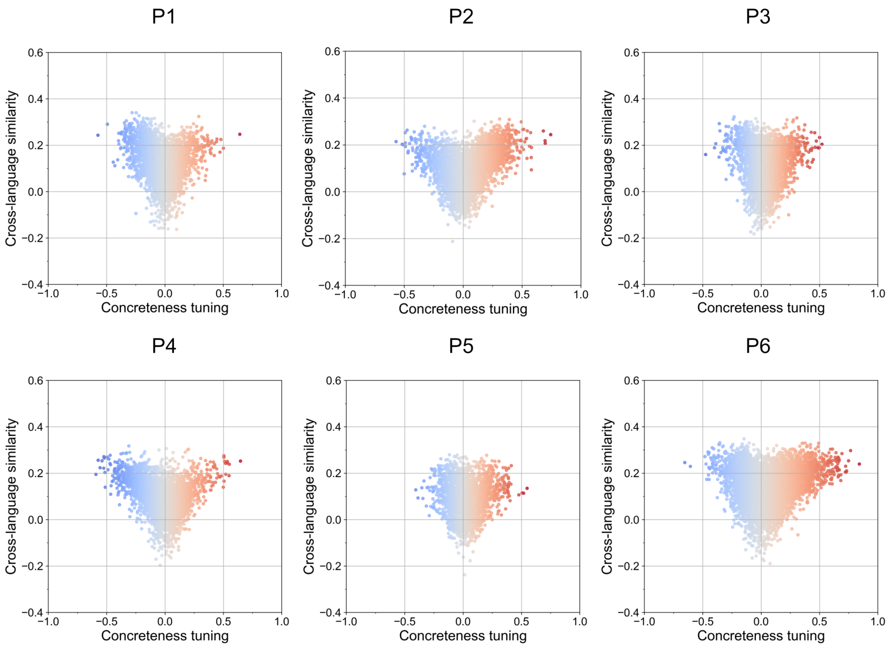


**Fig. S15 Comparison between concreteness tuning (x-axis) and cross-language similarity (y-axis) for all participants in multilingual BERT.** Each point represents a voxel that is well predicted in at least one language ($\sqrt{R_{en}^2} > 0.1$ or $\sqrt{R_{zh}^2} > 0.1$). Both voxels with very concrete or very abstract tuning have high cross-language similarity. This pattern reflects a positive correlation between the absolute value of concreteness tuning (towards concrete or abstract concepts) and cross-language similarity.

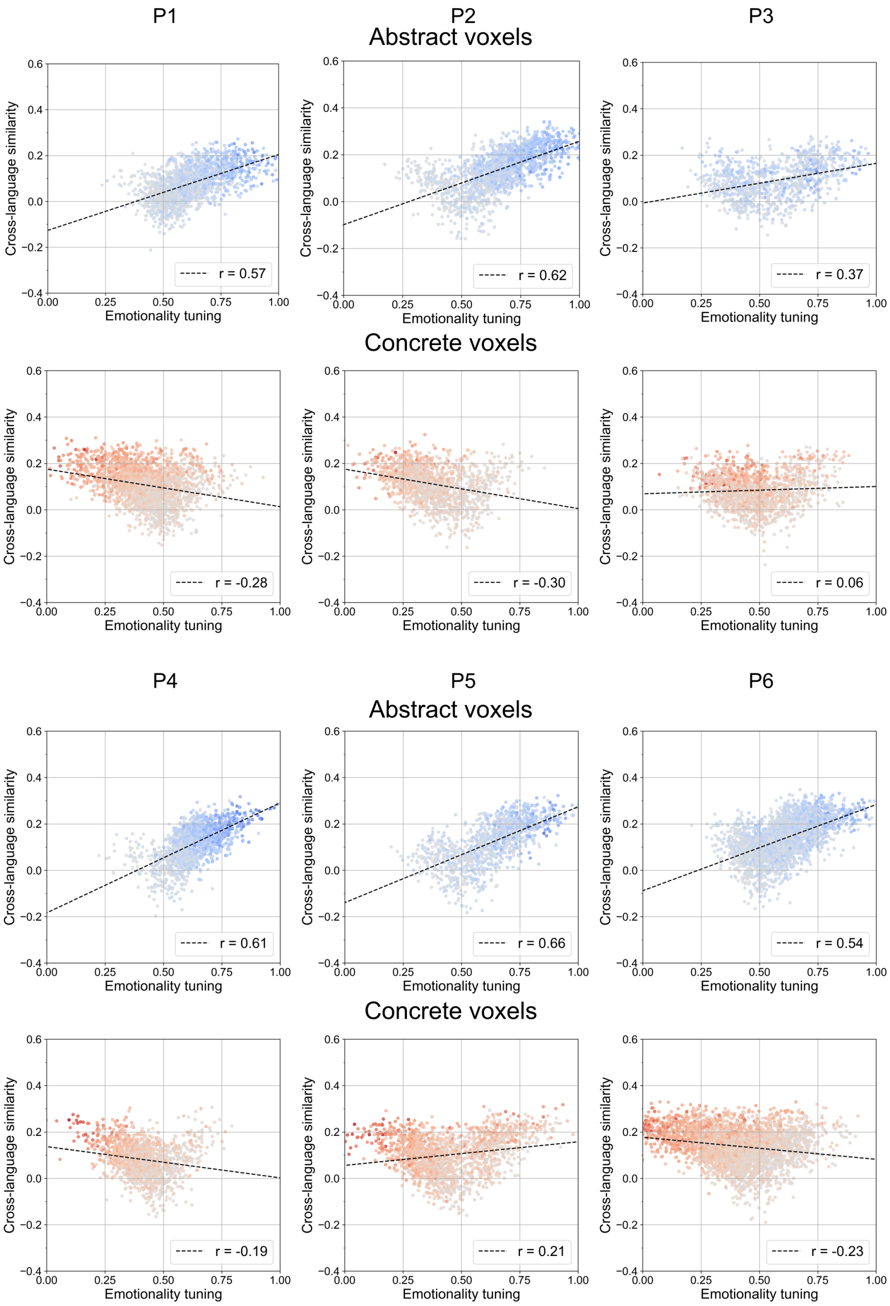


**Fig. S16 Comparison between emotionality tuning (x-axis) and cross-language similarity (y-axis) for all participants in multilingual BERT.** Voxels tuned towards abstract concepts (negative concreteness tuning) are shown on the top and voxels tuned towards concrete concepts (positive concreteness tuning) are shown on the bottom. Each point represents a voxel that is well predicted in at least one language ($\sqrt{R^2_{en}} > 0.1$ or $\sqrt{R^2_{zh}} > 0.1$). For all participants, the Pearson correlation between emotionality tuning and cross-language similarity (dotted line) is stronger for abstract-tuned voxels than for concrete-tuned voxels. This suggests that emotional content may drive cross-language similarity for abstract concepts.

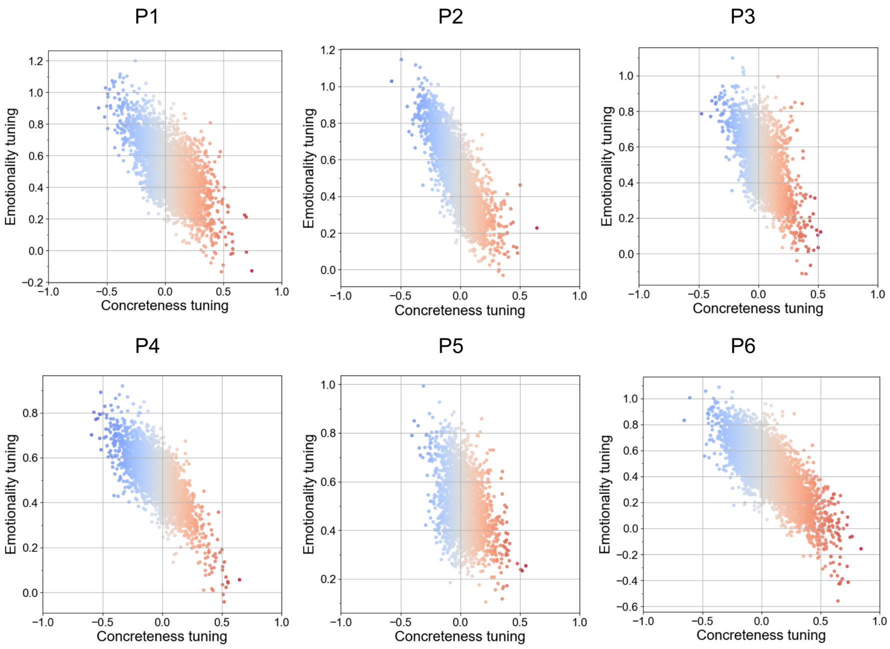


**Fig. S17 Comparison between concreteness tuning (x-axis) and emotionality tuning (y-axis) for all participants in multilingual BERT.** Each point represents a voxel that is well predicted in at least one language ($\sqrt{R^2_{en}} > 0.1$ or $\sqrt{R^2_{zh}} > 0.1$). There is a negative correlation between concreteness tuning and emotionality tuning. This suggests that regions tuned towards abstract concepts represent concepts with greater emotional content.

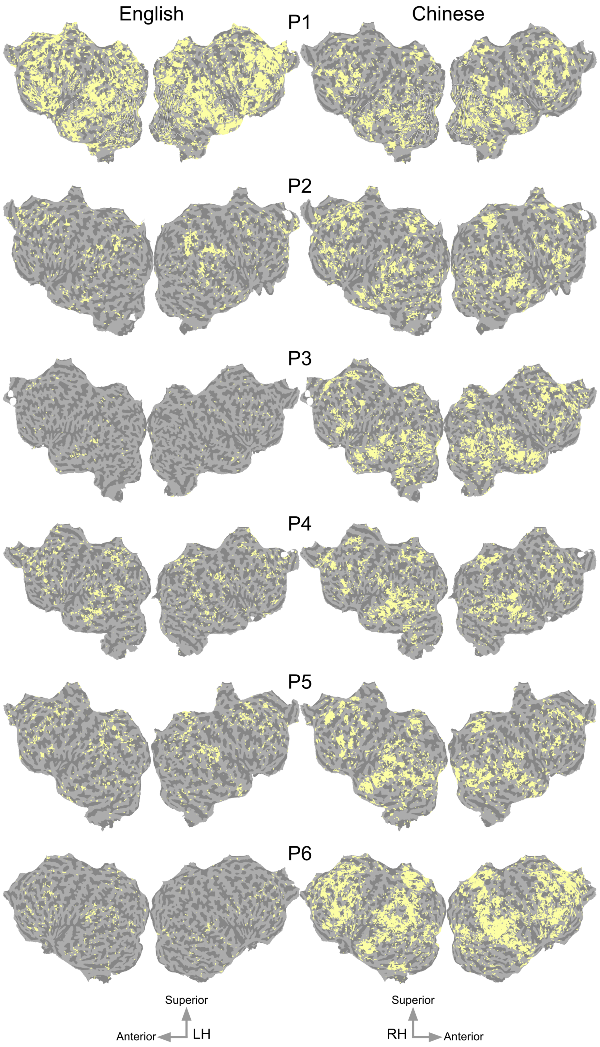


**Fig. S18 Statistical significance of prediction accuracy for each participant in each language in multilingual BERT.** The estimated semantic model weights of the multilingual BERT embedding space were used to predict voxel responses on a held-out story. Prediction accuracy was computed as the coefficient of determination ($R^2$) between predicted and recorded brain response on the held-out story. The statistical significance of prediction accuracy of each voxel was determined by comparing the prediction accuracy of the estimated models to the prediction accuracy in predicting permuted data. Voxels that were significantly predicted are shown on the flattened cortical surface of each participant, separately for each language. Voxels shown in yellow were significantly predicted (one-sided p¡.05, FDR corrected with a Benjamini-Hochberg correction for multiple comparisons).
